# Comparative genomic analysis reveals reduced pathogenicity of *Ralstonia* spp. in water

**DOI:** 10.1101/2024.10.23.619949

**Authors:** Gaopeng Liu, Chengzhi Mao, Qi Li, Da Huo, Tao Li

**Author notes:** Co-corresponding authors. Mailing addresses: Institute of Hydrobiology, Chinese Academy of Sciences, Wuhan 430072, Hubei Province, PR China. Corresponding authors E-mail: Qi Li Da Huo. **Conflict of Interest** The authors declare that they have no conflict of interest.

## Abstract

*Ralstonia* spp., known for their adaptability across various habitats, are known to cause infections. The adaptive metabolic diversity of those inhabiting aquatic ecosystems remains poorly understood. We report four new *Ralstonia pickettii* genomes enriched in the cyanobacterial culture derived from bloom-forming cyanobacteria, *Dolichospermum*. A total of 228 complete genomes from the *Ralstonia* genus were utilized for phylogenetic inference, categorizing them based on isolation environment and host: water, soil, plant, and human groups. Meanwhile, the abundance of carbohydrate-active enzymes and secondary metabolites in water and human groups differed from the plant-host associate habitat. *CeoB* and two β-lactamases types of *OXA* were identified in the water habitat, showing similarities to certain strains in the human-host but differences from other habitats. The infectivity within water habitats seems to diminish, as evidenced by the decreased abundance of T3SS virulence proteins. Moreover, a distinctive pyrimidine degradation pathway in water degrades exogenous pyrimidines to supply nitrogen and other compounds for energy metabolism to provide a potential for broader habitat adaptability. Fluorescence in situ hybridization results confirmed that *R. pickettii* rarely attached to cyanobacterial cells, indicating that they are not parasitic relationships. We postulate that the absence of T3SS and the unique metabolic profile represent adaptations of *Ralstonia* to an aquatic free-living lifestyle.

## 1. Introduction

*Ralstonia* spp. are Gram-negative non-fermentative bacteria within the family Burkholderiaceae, exhibiting wide distribution across diverse environments including water, plants, hospitals, and soil.^1^ Despite the widespread presence of *Ralstonia* spp. in several typical habitats, their occurrence in natural water bodies has received little attention compared with other habitats. Only a limited number of research reported that genomic islands (GIs) and horizontal gene transfer (HGT) are potential factors facilitating strong adaptability of *R. pickettii* in drinking water.^2^ More work needs to be done to uncover how *R. pickettii* adapts and survives in freshwater environments. Many species of *Ralstonia* spp. are opportunistic pathogens capable of infecting both humans and plants. In humans, they can lead to respiratory failure and potentially death, while in plants, it causes significant economic losses worldwide.^3,4^ Multiple studies have shown that *Ralstonia mannitolilytica* (*R. mannitolilytica*) and *Ralstonia pickettii* (*R. pickettii*, formerly known as *Burkholderia pickettii* or *Pseudomonas pickettii*) can cause severe infections in immunocompromised patients, leading to respiratory infections.^5,6^ *R. pickettii* also significantly impacts the safety of hospital water, capable of passing through 0.02 mm filter membranes and leading to contamination of sterile drugs.^7^ *R. solanacearum* can infect *solanaceous* crops, including tobacco, tomato, and potato, causing bacterial wilt and resulting in significant annual economic losses worldwide.^8,9^ Various factors contributing to *Ralstonia* spp. pathogenicity have been studied in *R. solanacearum*, including the type III secretion system (T3SS), exopolysaccharides (EPS), and flagella. However, the primary determinant of pathogenicity is the T3SS, which can inject type III effector proteins (T3Es) into the host cell cytosol to induce infection.^10^

Comparative genomics offers new insights into ecological niche adaptation within prokaryotes, such as the expansion of gene families in cyanobacteria that facilitate adaptation to terrestrial environments, and the identification of sulfur acquisition and metabolism as adaptive metabolic strategies in the genus *Novosibinobium*.^11,12^ As of now, a comprehensive view of phylogenetic relationships among species in *Ralstonia* genus remains unclear. Previous research has mainly focused on a specific group of species. Wicker et al. utilized multilocus sequence analysis (MLSA) to examine the intraspecific recombination patterns of *R. solanacearum*, categorizing them into four phylotypes corresponding to the geographic origins of the strains.^13^ Meanwhile, the *R. solanacearum* species complex (RSSC) is divided into *R. solanacearum* (phylotype II), *R. pseudosolanacearum* (phylotypes I and II), and *R. syzygii* (phylotype IV).^14,15^ For a long time, the genus of *Ralstonia* was considered comprising six species: *R. pickettii*, *R. solanacearum*, *R. pseudosolanacearum*, *R. syzygii*, *R. mannitoliytica*, and *R. insidiosa*. Recently, Lu et al. reported three novel species isolated from tobacco fields in Yunnan, China, forming an independent clade.^16^ *R. solanacearum* phylotype I represents one of the ongoing diversifying subspecies, spreading from the lowlands to the highlands in China.^17^ The rapidly increasing number of complete genomes in databases, with over 600 genomes of *Ralstonia* spp. available in the National Center for Biotechnology Information (NCBI) database, offers unique opportunities to study the evolutionary relationships among these organisms.

In our recent survey of a *Dolichospermum* bloom event, known for producing cyanotoxins such as saxitoxin and anatoxin-a^18^, we isolated cyanobacterial filaments from a lake in Wuhan, China. The metagenome of the culture shows a significant number of *R. pickettii* coexisting with *Dolichospermum* cells. We recovered four complete genomes from the metagenomes. By conducting a comparative genomic analysis, including phylogenetic analysis (*Ralstonia* spp. categorized into four distinct clades based on habitat and host conditions: soil, water, plant and human groups), analysis of antibiotics resistance genes, CAZyme analysis, exploration of secretion systems, secondary metabolite and metabolic analysis of common and shared features among different clades. Our objective is to elucidate the adaptation mechanisms of *R. pickettii* to water environments, explore the pathogenic changes of *Ralstonia* spp. in various habitats, and investigate the potential relationship between *R. pickettii* and *Dolichospermum* blooms.

## 2. Materials and methods

### 2.1 Sampling and identification

On May 17th, 2022, a water sample was collected from a natural lake (N: 30.546166, E: 114.353721) in Wuhan, Hubei Province, China. Algae filaments were separated, washed multiple times with distilled water droplets, and then cultured in CT medium. Bacterial identification was conducted using the online BLAST program (http://blast.ncbi.nlm.nih.gov/Blast.cgi). The ncbi-genome-download (v 0.3.3) (https://github.com/kblin/ncbi-genome-download) tool was employed to download all genome sequences of the *Ralstonia* genus from the RefSeq section, and pyani (v 0.2.12)^19^ was used to calculate the average nucleotide identity (ANI) for further species identification under -m ANIm -g parameter. The culture used for sequencing was maintained at 28 ± 1°C with a speed of 100 rpm in the incubator.

### 2.2 Genome sequencing, assembly, and annotation

Metagenomic DNA samples obtained from culture were extracted using the DNA Gel Extraction kit according to the manufacturer’s instructions (Axygen, USA). DNA concentration was measured using a Nanodrop 2000 spectrophotometer (Thermo Scientific, USA). Library construction and sequencing were conducted at BENAGEN Biotechnology Co., Ltd. Sequencing was performed using the Illumina MI seq PE150 platform for next-generation sequencing and the Nanopore PromethION platform for third-generation sequencing, generating 5 Gb of data, respectively. Offline data processing was performed using the fast Guppy basecaller (v 6.3.8) for accurate base calling with a high-accuracy model and subsequent quality control. FastQC (v 0.11.9)^20^ was used for quality control, and raw data with a quality score below Q20 was discarded. Filtering sequences shorter than 5000 bp, de novo assembly was conducted using the Flye (v 2.9.1) - Racon (v 1.5.0) model^21,22^ with nanopore sequences under default parameter, followed by three rounds of correction and polishing. Next, Illumina sequencing data was used for two rounds of polishing of the de novo genome using NextPolish (v 1.4.1)^23^ to eliminate sequencing errors under default parameter. *R. pickettii* genomes data were extracted from the complete metagenomic dataset and annotated using Prokka (v 1.14.6) under default parameter.^24^ Full-length 16s rRNA sequences were extracted from the genome using barrnap (v 0.9) (https://github.com/tseemann/barrnap) software and identified using the online BLAST program (http://blast.ncbi.nlm.nih.gov/Blast.cgi).

### 2.3 Phylogenetic analysis

For comparative analysis, we retrieved 428 *Ralstonia* spp. genome sequences and their annotations from the NCBI GenBank database. We selected 228 genomes with completeness above 99% (evaluated using CheckM (v 1.2.1)^25^ under lineage_wf mode and fewer than 100 contigs for further analysis. Accession numbers of the genomes included in this study and their genomic features are shown in (Supplementary Table S1). Protein sequences were extracted from GenBank files using a custom python script, and single-copy orthologous gene families were identified for the 231 genomes (including three species selected as outgroups) using OrthoFinder (v 2.5.5) under -a 50 -M msa parameters.^26^ We identified 421 single-copy orthologous proteins with an average length of more than 100 bp for further analysis. After alignment and trimming using MAFFT (v 7.520)^27^ under --auto parameters and TrimAl (v 1.4) under -automated1 parameters,^28^ respectively, conserved proteins for each species were concatenated using SeqKit (v 2.5.1).^29^ The concatenated amino acid sequence of the 421 gene families was used to construct a maximum likelihood (ML) phylogenomic tree with ultrafast 1000 bootstrap replicates using iqtree2 (v 2.2.5) under -m MFP -B 1000 -bnni parameters.^30^ The amino acid substitution model was automatically selected the best-fit model suggest by iqtree2 software. Phylogenetic trees were visualized and beautified using iTOL (v 5.0)^31^ and Adobe Illustrator CC 2019.

### 2.4 FISH and microscopy

The cultures in logarithmic growth phase were centrifuged at 8000 rpm for 10 minutes, washed three times with sterile PBS (phosphate buffered saline), and fixed overnight at 4 □ in 4% paraformaldehyde. Glass slides were pretreated with 0.1% polylysine (Sangon Biotech), onto which the fixed cultures were added and dried at 45 □ for more than 20 minutes to allow adsorption onto the slides. After fixation, pre-hybridization was performed using a hybridization solution and incubated at 40 □ for 1 hour. Subsequently, a 30 μl solution of probe (5‘-GCAAGGCCTCATGCTATAG-3’, diluted 1:5 v:v in 30% formamide) was added and incubated overnight at 40 □ in a moist chamber. Washing steps included 15-minute washes with 2 × SSC (Saline sodium citrate), followed by two 7 minutes washes with 1 × SSC, 15 minutes wash with 0.5 × SSC, and 45 minutes incubation with diluted branch probe at 40 □ in a moist chamber, followed by 7 minutes washes with 2 × SSC, 1 × SSC, 0.5 × SSC, and 0.1 × SSC. Incubation at 42 □ for 3 hours with a fluorescent signal probe in a moist chamber was followed by the same washing process. DAPI solution (2 μg/ml) was incubated for 20 minutes away from light, and the samples were gently rinsed three times with sterile PBS before sealing the slides with an anti-fluorescence quenching agent. The dilution ratio of SSC solution is expressed as volume ratio (v/v). All reagents were obtained from Wuhan Servicebio Biotechnology Co., Ltd. The design and synthesis of *R. pickettii*’s fluorescent probe were completed by Wuhan Servicebio Biotechnology Co., Ltd. Observations and panoramic scanning were conducted using a Leica microscope Aperio VERSA 8 (v 1.4.0.125). DAPI staining was observed using ultraviolet excitation luminescence, *Dolichospermum* sp. self-luminescence was observed using an excitation wavelength of 510-560 nm (G-2A), and *R. pickettii* was observed emitting green fluorescence in the wavelength range of 460-500 nm.

### 2.5 Comparative genomics analysis

Gene function and secretion system prediction were conducted using KofamScan (v 1.3.0)^32^ with database version 20231127 under default parameter, and the software recommended annotation was selected as the final result. CAZymes annotation was performed using dbCAN2 (v 4.0.0)^33^ against the dbCAN2 database (v12) with hmmer-based comparison (coverage > 0.35 and e-value < 1e-15), dbCAN-sub (coverage > 0.35 and e-value < 1e-15), and diamond (e-value < 1e-10) methods. The annotation predicted by hmmer was used as the statistical standard after integrating the results from three methods. Antibiotic resistance genes (ARGs) and virulence factors were predicted using abricate (v 1.0.1) (https://github.com/tseemann/abricate) with the card^34^ and VFDB^35^ databases, respectively (identity > 80%). Bacterial secretion systems prediction was extracted from KofamScan annotated results, and gene names and classification information can be queried through KO numbers. Annotate secondary metabolites using online and detection strictness used strict, and other parameters using default methods (https://antismash.secondarymetabolites.org/#!/start).^36^

A python script was employed to screen OGs (orthogroups genes, OGs) present in all strains, based on the orthofinder results excluding outgroups, shared metabolism representing extremely conservative sequences that present in all genomes. The incomplete transporter with one lost and aligning it with the corresponding protein sequence of *E. coli* in BLASTp (identify>50%). To investigate unique metabolic content in each group, a precise Fisher test (p < 0.05) was conducted to confirm whether certain metabolic pathways were lost in specific groups, thus obtaining unique metabolic content for each group. The longest aligned sequence representing each selected OG was chosen, and all sequences were merged for annotation using KofamScan (v 1.3.0)^32^ with database version 20231127. The annotation table was reconstructed using KEGG-reconstruct (https://www.kegg.jp/kegg/mapper/reconstruct.html) to visualize metabolic pathways. A simplified metabolic diagram was created using Adobe Illustrator CC 2019 based on the metabolic pathway results. *Ralstonia* genus used Badirate (v 1.35) software^37^ based on BDI-CSP-FR model to make gene family contraction and expansion which provide an important perspective of species evolution. The standard binary phylogenetic tree, which has removed duplicate values using for gene family contraction and expansion analysis.

### 2.6 Statistical analysis and data visaliization

Data were analyzed and visualized using R (v 4.3.1) in RStudio (v 2023.6.0.421) with the R packages pheatmap (v 1.0.12) and ggplot2 (v 2.3.4). Results for CAZymes, resistance genes, and virulence factors were generated using pheatmap (v 1.0.12) and gplots (v 4.3.1). Boxplots for genome-related information were created using ggplot2. The vegan package (v 2.6.4) was utilized for analysis of similarities (anosim) to distinguish among different clades based on CAZymes.

## 3. Results

### 3.1 Phylogenetic reconstruction

We selected 228 genomes with completeness above 99% and fewer than 100 contigs for further analysis. The ANI results of the 228 genomes (excluding 3 outgroups) (Supplementary Fig. S1) indicate that the similarity between the four *Ralstonia* strains and *R. pickettii* is over 98 %. Based on the primary habitat of the strains within each clade, the 228 species can be categorized into four groups: plant, human, soil and water (Fig. 1A). In the plant group, there are *R. solanacearum*, *R. pseudosolanacearum* and *R. syzygii*. There are *R. insidiosa* and *R. mannitolilytica* in the human group, novel species are found in the soil group and *R. pickettii* in the water group. The average genome length and GC content of these 228 *Ralstonia* spp. genomes were 5.49 Mb and 64.99 %, ranging from 4.38-7.02 Mb and 62.95-67.1 %, respectively (Fig. 1B, 1C). The greatest variation is the GC content across different habitats, clades associated with plant-hosts comprising the highest proportion, averaging 66.61%, while soil group exhibits the lowest average content at 63.60%. The GC content related to groups of plant and human are significantly higher than in other habitats, indicating stronger genomic stability in these habitats. However, the number of CDS showed a significant increase in soil and water habitats compared with the plant group (Fig. 1D). In this study, it was observed that the number of gene families in the common ancestor of the *Ralstonia* genus was relatively modest, with an increase in gene family count occurring concomitant with adaptation to diverse environments. The *R. insidiosa* gene family, which is associated with human, boasts the highest number of genes and may be implicated in its adaptation to various hosts or environment (Supplementary Fig.S2).

**Fig. 1.**
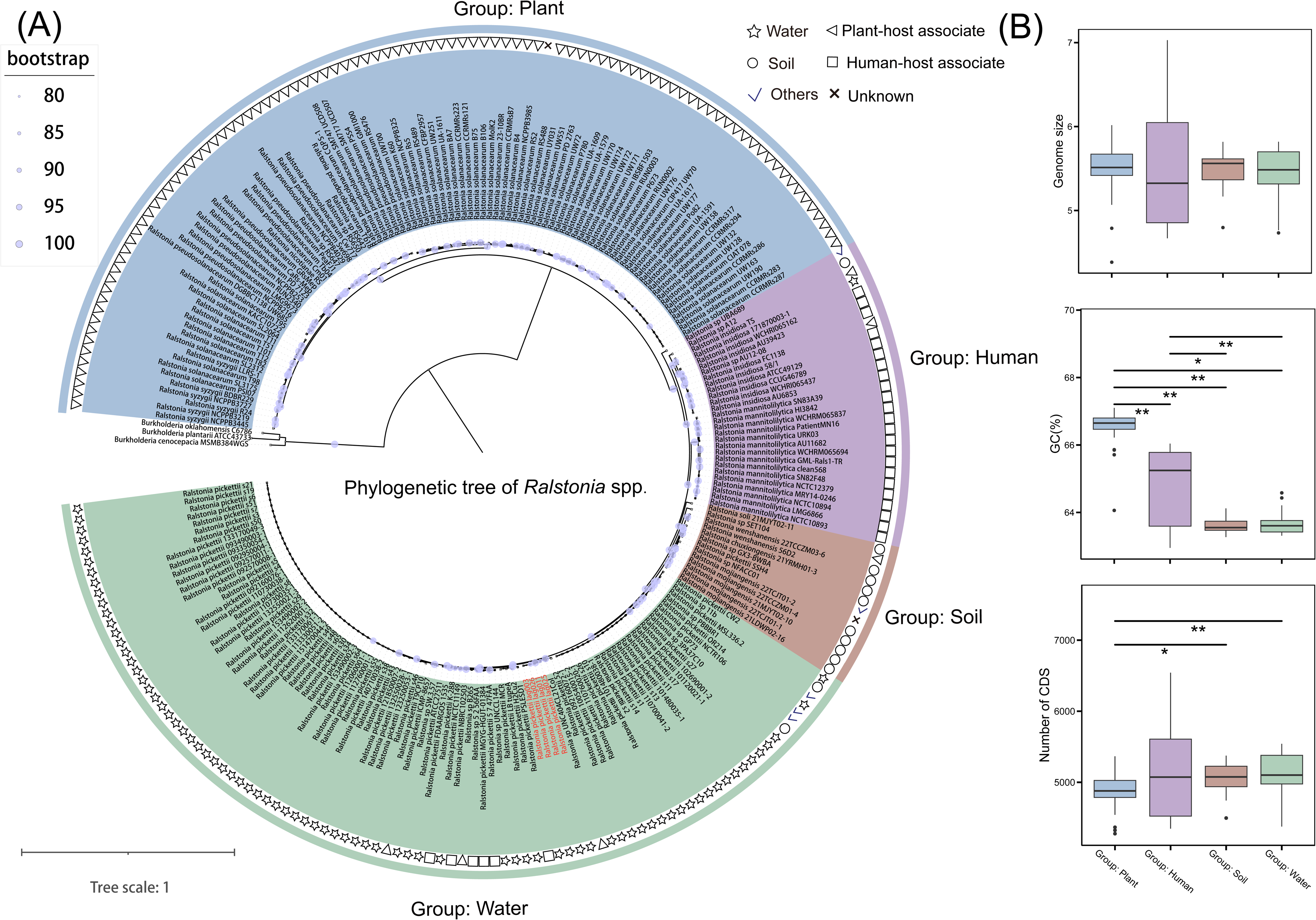
Information statistics and phylogenetic reconstruction of *Ralstonia* spp. genomes. (**A)** Phylogenetic analysis of the *Ralstonia* spp. genomes. The maximum likelihood tree was constructed based on the concatenated alignment of 421 single copy orthologous proteins. Nodes with bootstrap value ≥ 50 are indicated by solid circles. (**B)** Genome size, GC (%) content and number of CDS distribution in different clades. The significant differences between pairs were determined using the Wilcoxon test, with significance indicated by * (*p* < 0.05) and ** (*p* < 0.01).

### 3.2 Type of toxicity and infectivity in Ralstonia spp

ARGs are present in nearly all genomes and can be divided into three types: *OXA*, *ceoB*, and *sul2* (Fig. 2). In the plant group, *ceoB* is present in almost all genomes, and it is related to a cytoplasmic membrane component of the CeoAB-OpcM efflux pump, which effluxes antibiotics. In the human group, the *OXA* only present in *R. mannitolilytica* and *ceoB* present in all strains. These belong to the *OXA* beta-lactamase family and have the ability to inactivate antibiotics. The water group contains *OXA* and *ceoB* in almost every genome, with only one strain having *sul2*, a sulfonamide-resistant dihydropteroate synthase of Gram-negative bacteria usually found on small plasmids, which can replace the antibiotic target. The virulence factor *bopC* is exclusively distributed in soil group, inducing host cell necrosis. *FlgG*, a common virulence factor activating the host’s innate immunity, is found in some groups of plant and human genomes.

The bacterial secretion system plays a crucial role in the invasion of pathogenic microorganisms into hosts and serves as an indicator of toxicity. All genomes in this study contain complete T2SS, the typical T2SS is a secretion system that transports substances from the periplasmic space to the outer membrane of bacteria, encoded by approximately 12 to 15 *gsp* (general secretion pathway) genes, indicating a potential infection ability in *Ralstonia* spp. T3Es secreted by T3SS play a pivotal role in pathogenesis. Interestingly, the number of T3SS proteins is significantly lower in water group compared to other groups. While T4SS proteins is present in all clades, but with a higher presence observed in water group. Conversely, T6SS exhibits a certain degree of deficiency in water group.

**Fig. 2.**
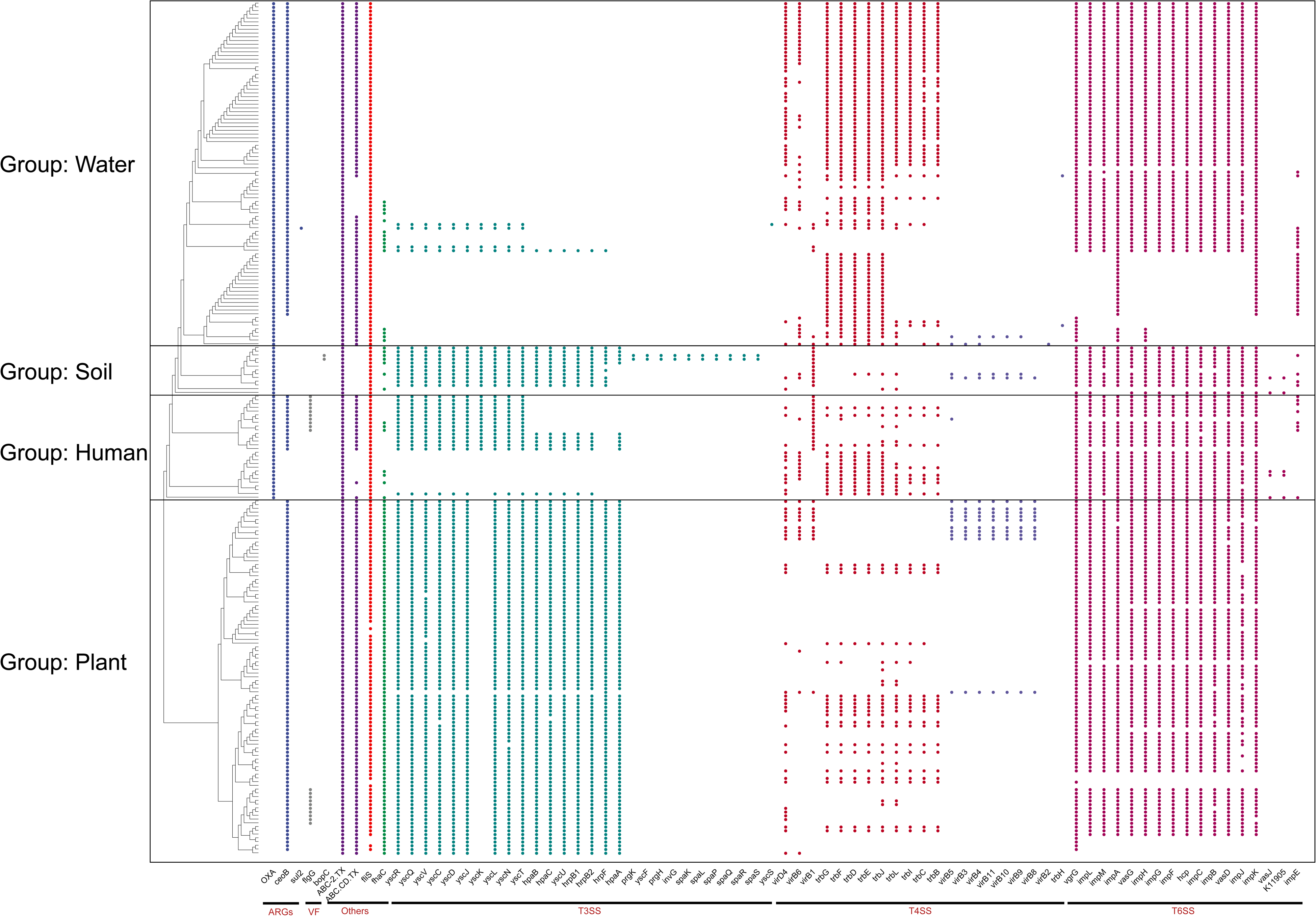
Analysis of antibiotics resistance genes (ARGs) and bacterial secretion system composition in *Ralstonia* spp. genomes. **A** ARGs and virulence factors within the genome of 228 *Ralstonia* spp., *OXA-22, OXA-443*, *OXA-444*, *OXA-60*, *ceoB* and *sul2* are ARGs, *bopC* and *flgG* are virulence factors. **B** Diversity analysis of different bacterial secretion systems in *Ralstonia* spp. genomes. T3SS (type III secretion system), T4SS (type IV secretion system), T6SS (type VI secretion system), Others (Other secretion system), T2SS (type II secretion system) is complete in all groups and is not shown in the figure.

### 3.3 CAZymes in Ralstonia genus

CAZymes are involved in biological processes related to carbohydrate synthesis and metabolism. They play roles in synthesis (GTs), degradation (GHs, PLs, CEs, AAs), and recognition (CBMs). The proportion of GHs, involved in carbohydrate degradation, was more abundant in the plant group compared to other groups (Fig. 3A). Conversely, there were more GTs in the human group, which may be related to host invasion. There were no significant differences observed among PLs, CEs, CBMs, and AAs across the groups. The group of water species contains a different abundance GTs, indicating that different water habitat can also affect the composition of CAZymes. NMDS results revealed differences between the plant group and those groups with other habitats (R=0.776, *p*=0.001) based on non-parametric Bray-Curtis distance (Fig. 3B).

**Fig. 3.**
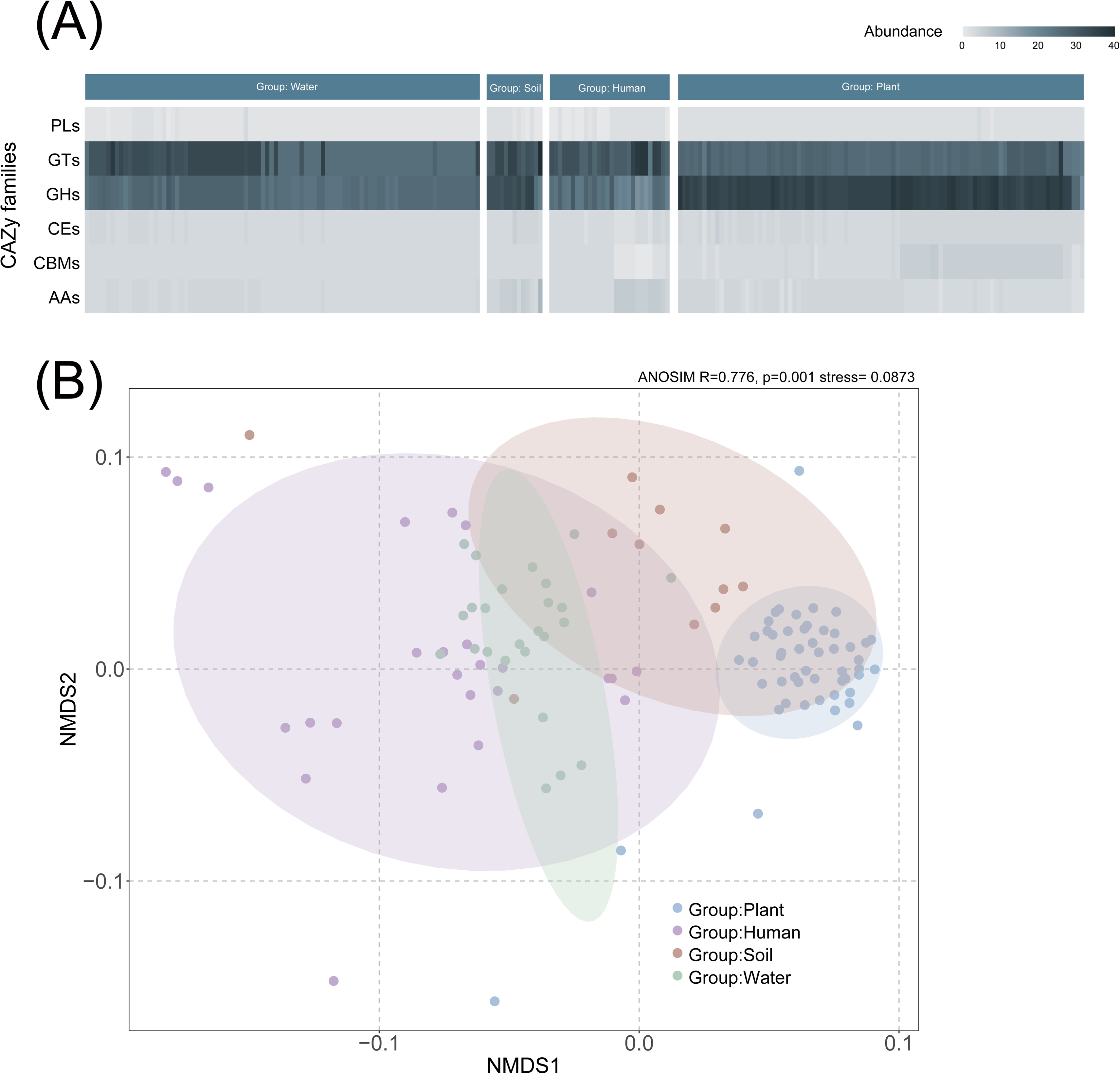
Carbohydrate active enzymes (CAZymes) composition and difference analysis in *Ralstonia* spp. genomes. **A** NMDS (Non-metric multidimensional scaling, NMDS) analysis of different clades composed of enzymes based on Bray-curtis distance. **B** Diversity and abundance of CAZymes in different clades. CAZyme genes are classified into PL (polysaccharide lyase), GT (glycosyltransferase), GH (glycoside hydrolase), CE (carbohydrate esterase), CBM (carbohydrate-binding module) and AA (auxiliary activity) categories.

### 3.4 Metabolism analysis of Ralstonia spp

The metabolism of microorganisms is highly diverse, and the core genome of the *Ralstonia* spp. consists of only 12 pathways or biochemical processes, including amino acid metabolism (leucine biosynthesis, proline biosynthesis and degradation), energy metabolism (cytochrome-c oxidase and oxygen oxidoreductase), metabolism of cofactors and vitamins (lipoic acid biosynthesis), carbohydrate metabolism (glycolysis of three-carbon compounds, nucleotide sugar biosynthesis, ribose-phosphate pyrophosphokinase), and nucleotide metabolism (adenine ribonucleotide biosynthesis, guanine ribonucleotide biosynthesis, ATP phosphohydrolase). There are 3 shared ABC transporters, including absorption of branched-chain amino acids, D-methionine, and efflux of LPS (lipopolysaccharide, LPS). Results of dismissd transporters in whole genome searches using BLASTp tools based on *E. coli* sequences (identity > 50%) indicate incomplete transporters and loss of one protein (Fig. 4).

The phthalate degradation and acetate synthesis pathways are observed in plant and water groups, while phenol 2-monooxygenase is absent in soil group. The pyrimidine degradation pathway is only present in the group of water, while PEP (phosphoenolpyruvate) synthesis is present in plant and water groups. There are 4 types of ABC transporters observed with dismiss based on BLASTp result in all genomes, including Fe^3+^, phosphate, phosphonate and glutamate/aspartate.

**Fig. 4.**
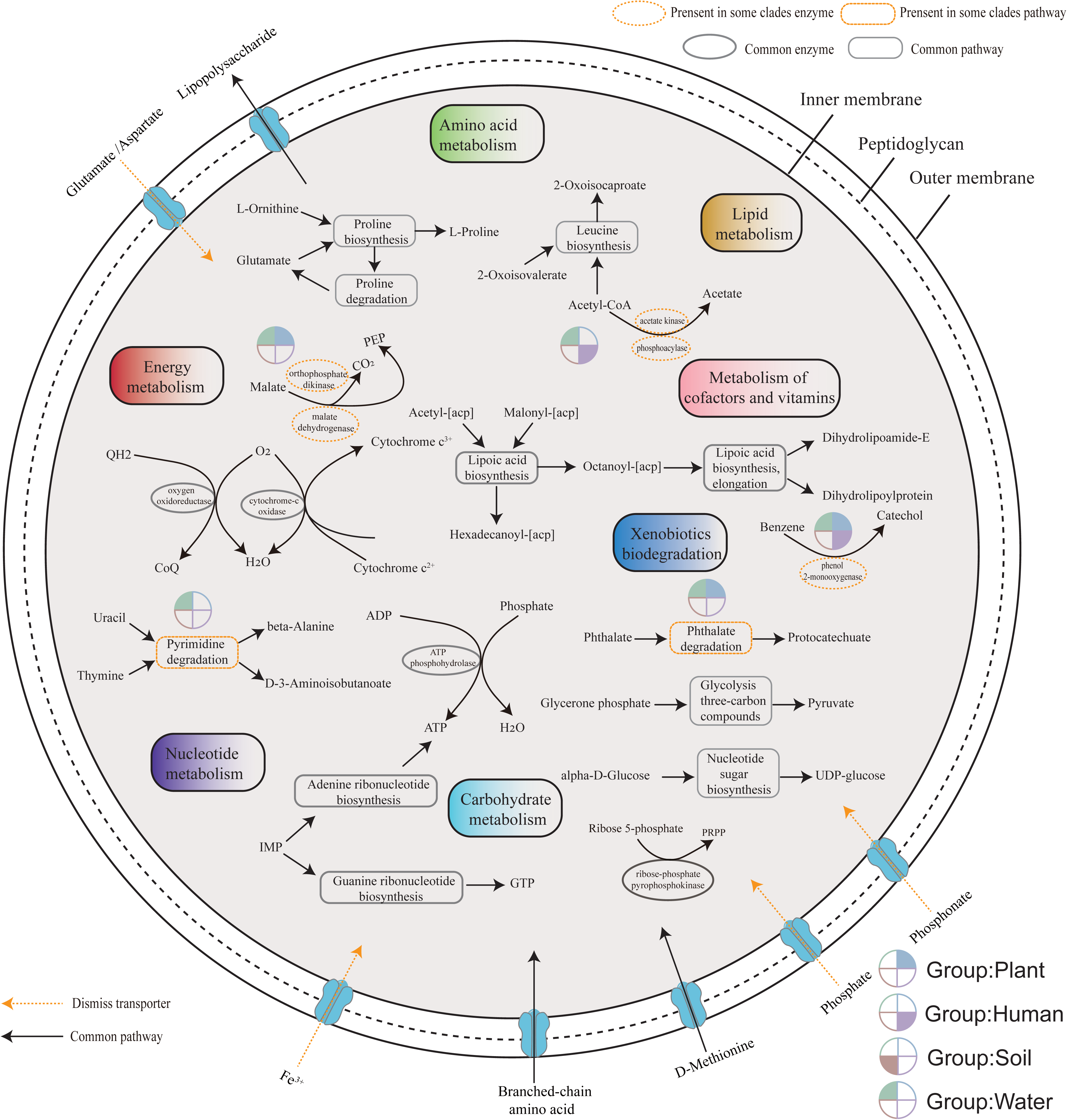
Overview of the metabolic potential of different habitats. **A** Metabolic pathways were constructed based on the orthogroups genes count matrix (excluding outgroups) from OrthoFinder results, the longest aligned orthogroups genes’ amino acid sequence was selected as the representative sequence for KEGG annotation. Genomic shared metabolism representing extremely conservative sequences that present in all genomes. **B** Metabolic analysis dimiss to each clade, which based on Fisher’s test (*p* < 0.05). The incomplete transporter with one lost and aligning them with the corresponding protein sequences of *E. coli* in BLASTp (identify>50%).

### 3.5 Secondary metabolite prediction from Ralstonia spp. genomes

The prediction of secondary metabolites in *Ralstonia* spp. using antiSMASH revealed a total of 16 secondary metabolites, with varying proportions observed across different habitats (Fig. 5A). The average secondary metabolite content of each genome in different habitat is highest in the plant group, with NRPS (Nonribosomal peptides, NRPS) content being the highest. However, in the human and soil groups, the average content of arylpolyene is the highest, while in water group, the content of redox-cofactor is the highest. Interestingly, the content of redox-cofactor in the plant group is extremely low, while the content of T1PKS is higher than the others. At the same time, the results of NMDS also indicate differences in secondary metabolites between plant group and other groups (stress value < 0.2, R=0.545, *p*=0.001) (Fig. 5B).

**Fig. 5.**
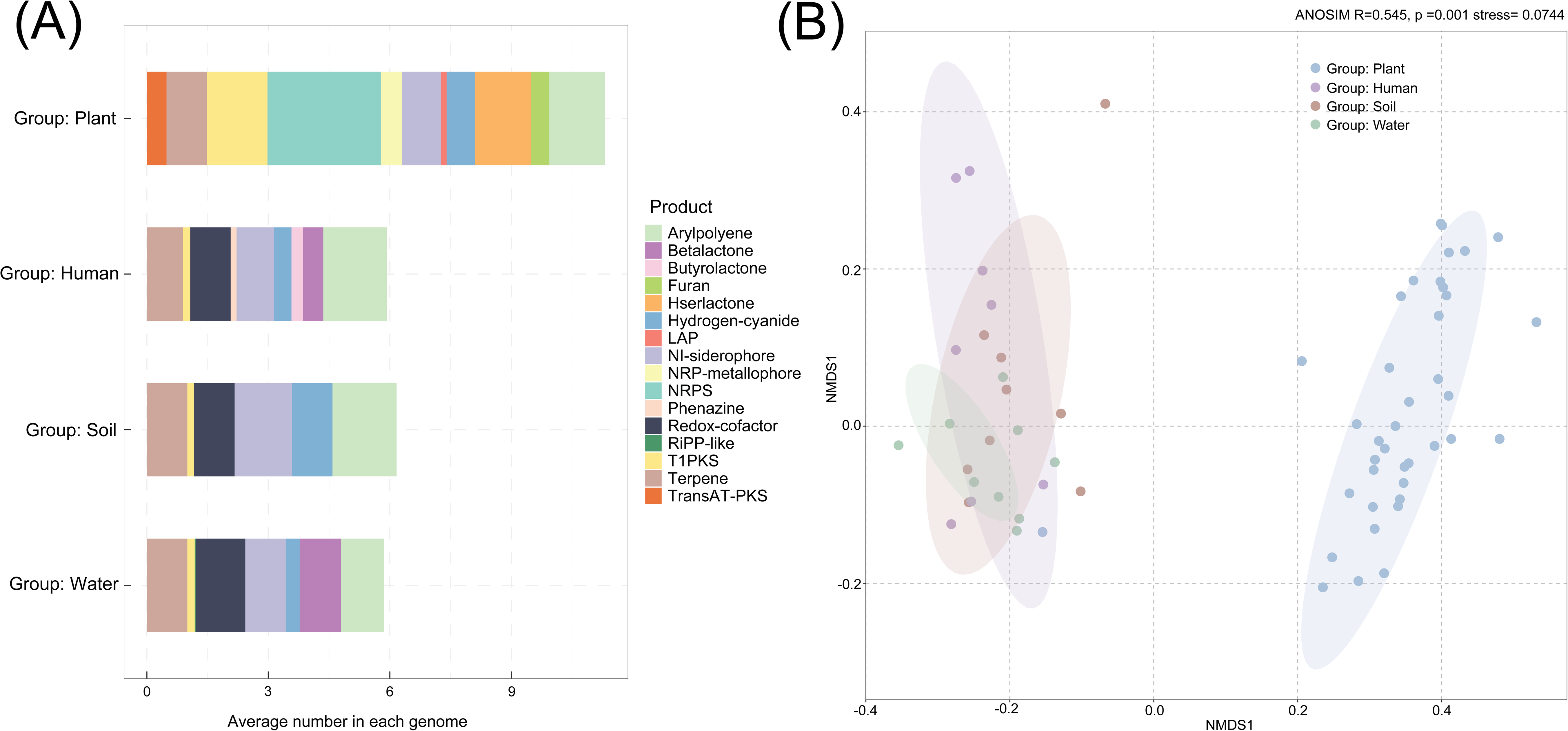
Prediction of secondary metabolites of *Ralstonia* spp. in different habitats by antiSMASH analysis. **A** The average amount of secondary metabolites contained in each genome in different habitats. **B** NMDS analysis of different clades composed of secondary metabolites based on Bray-curtis distance.

### 3.6 Localize R. pickettii in Dolichospermum culture

Using fluorescence in situ hybridization (FISH) to analyze the relative positions of *Dolichospermum* sp. and *R. picketti* revealed significant aggregation, with a notably higher abundance of *R. picketti* colonies observed in the field (Fig. 6A-C). The morphology of *Dolichospermum* filaments is distinct, emitting red excitation light *R*. *pickettii* rarely attaches on the *Dolichospermum* cells. However, there is a small amount of aggregation of *R. pickettii* within the EPS blocks surrounding the filaments. The CT medium is relatively carbon deficient, thus, the *Dolichospermum* sp. may serve as carbon producers within microbial communities by generating a substantial number of EPS.

**Fig. 6.**
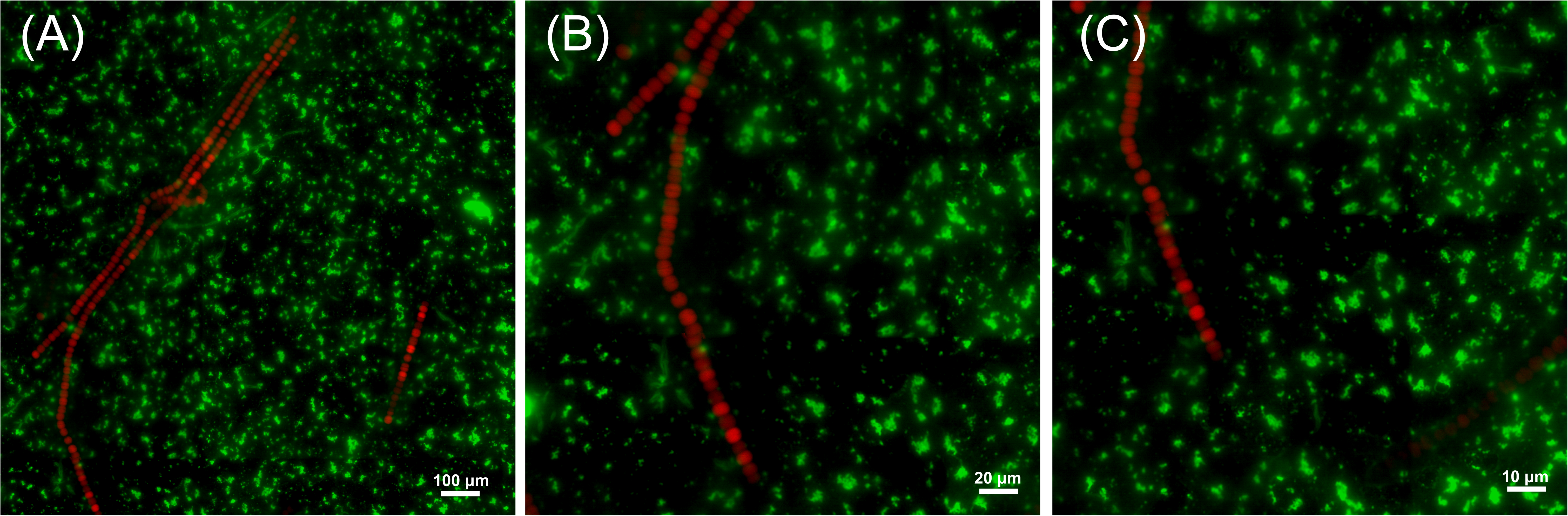
The relative positional relationship between *R. pickettii* and *Dolichospermum* sp. in the culture medium. Using a probe designed based on the 16s rRNA sequence of *R. pickettii*, light is activated within the 460-500 nm wavelength range (green), whereas the self-luminescence of blue-green algae (red) is activated within the 510-560 nm (G-2A) wavelength range. Maximum intensity projection of stack fluorescence images acquired at magnifications of ×10 (**A**), ×20 (**B**), and ×100 (**C**) is shown.

## 4. Discussion

Our comparative genomic analysis utilized single-molecule sequencing to assess pathogenicity and metabolic dissimilarity among *Ralstonia* species, with particular attention to the aquatic habitat lineage. The position of the evolution herein named *Ralstonia* has been unclear with environment and host. Phylogenomic trees are essential for a meaningful interpretation of the biological and evolutionary relationships of all organisms.^38^ Here, the main habits are divided into four clades clearly, namely plant, human, soil and water, the evolutionary history is closely correlated with environmental selection. The group related to plant-host is clearly distinguished from the other clades. The clustering of the other clades suggests a potential inconsistent habitat and evolutionary status of species among them due to human activities.

### 4.1 T3SS deficiency and ARGs diversity increased in water group

Many *Ralstonia* species are opportunistic pathogens that infect humans and plants, causing respiratory failure and leading to significant economic losses, respectively.^3,4^ *R. pickettii* in water can pass through the 0.02 mm filter membrane and contaminate physiological saline, causing an outbreak of leukemia that lasts for two weeks.^7^ *R. pickettii* is highly pathogenic in water, it will have a great impact on the safety of drinking water during algal blooms. Here, we predict the virulence effects of *Ralstonia* genus through analysis of ARGs, virulence factors, and infection systems. The production of antibiotics by microbes (especially in fungi) is a mode for them to gain advantages in microbial communities.^39^ Meanwhile, bacteria have evolved various strategies to resist antibiotics, such as enzymatic inactivation, target alteration, efflux, and permeability changes. OXA can activate the antibiotic inactivation method to protect cells from harm by antibiotics.^40^ The presence of different isomers (*OXA-22*, *443*, *444*, *60*) in the *Ralstonia* genus may extensively harbor this antibiotic exclude plant group, highlighting the necessity for vigilance regarding antibiotic resistance in water, soil, and human-hosts groups. *CeoB*, a cytoplasmic membrane component of the CeoAB-OpcM efflux pump, possesses the ability to pump antibiotics out of a cell, conferring resistance to bacteria.^41^ There are multiple *OXA* isomers and *ceoB* in water, at the same time, the similarity of ARGs composition among the three habitats except the plant-host clade. The reasons for the diversity of ARGs are closely related to the microenvironment, where ARGs can be transferred among various microorganisms through mobile genetic elements (MGEs), including transposons, plasmids, and insertion sequences, thereby facilitating the adaptive evolution of resistant bacteria in this environment.^42,43^

Gram-negative bacteria possess six types of secretion systems, labeled from I to VI. *R. solanacearum* relies on the TSS for the production of EPS, cell appendages, and protein secretions.^44^ In the plant host-associated clade, the T3SS, renowned for its direct transmembrane transport mechanism, has been observed to secrete a diverse array of T3Es. Some studies have reported the secretion of up to 74 T3Es, including PopA, PopB, and PopC.^45^ These proteins play a critical role as essential prerequisites for bacterial invasion and colonization of plant vascular systems.^9^ T2SS and T6SS have the capability to transport proteins in both fully or partially folded states, whereas other systems predominantly transport unfolded proteins. T2SS is not a complete transmembrane secretion system; it depends on the SecYEG (Sec export system) or Tat export system (twin arginine translation) to transport substances to the periplasmic space before being transported to the extracellular space. Many pathogens utilize this secretion system to secrete proteins, for instance, human pathogens like *Vibrio cholerae*,^46^ and *Aeromonas hydrophila*, which can infect humans, livestock, and aquatic animals.^47^ T2SS-dependent plant cell wall degrading enzymes in *R. solanacearum* confer advantages to invasive plants.^48^ Plant pathogens such as *Dickeya dadantii* utilize T2SS to infect the host.^49^Although the infection ability of *R. pickettii* in water is weaker than in other habitats in terms of the number of T3SS, *R. pickettii* may pose a threat to human health through other unknown mechanisms. The diversity of *R. pickettii* in global freshwater still deserves the attention to further research.

### 4.2 CAZymes and secondary metabolites different between plant group and others

Additionally, we explored the survival patterns of different groups in the habits through CAZymes differences. CAZymes encompass all enzymes involved in biological transformations related to sugar synthesis and metabolism. Additionally, CAZymes exhibit a higher prevalence in the genomes of pathogens, potentially associated with the degradation of complex carbohydrate structures within hosts, such as plant cell walls.^50^ Plant-host group is distinguished from other groups, the abundance of GHs associated with the plant-host was found to be higher compared to other clades in this study, these may be related to the composition of the invading plant’s cell walls. GTs are closely associated with protein glycosylation in pathogenic bacteria, influencing their virulence by promoting adhesion to host cells.^51^ Currently, GTs have been implicated in various pathogenic bacteria, including enterotoxigenic *E. coli*, *Photorhabdus asymbiotica*, *Pseudomonas aeruginosa*.^52,53^ In this study, the abundance of GTs showed variation even in the water group, potentially due to differences in aquatic environments.

Secondary metabolites are substances that use primary metabolites as precursors and do not have specific biological functions. Due to different hosts, there are significant differences between plant-host associate and other groups, such as NRPS, which is a diverse peptide substance that plays an important role as an antibiotic in clinical use.^54^ This plays an important role in enhancing the competitiveness of endosymbiotic microbial communities in plants. The proportion of redox-cofactor and betalactone are the highest in aquatic environments than other habitats. Redox cofactor are molecules that contains multiple types and can enhance the ability of microorganisms to adapt to various extreme environments.^55^ Betalactone compounds contain multiple antibiotics and are an important way to obtain dominant community ecological niches.^56^ The results indicate that *Ralstonia* spp. has adapted to a variable environment and produced different types of secondary metabolites.

### 4.3 A distinctive pyridine degradation pathway in the water group

The metabolism of microorganisms is diverse, consists of only 12 pathways or biochemical processes represents an extremely conservative function under strict conditions (present in all genomes). In the water and human groups, acetate metabolism is complete which an excessive organic acids can inhibit the growth of *E. coli*,^57^ a characteristic utilized in preventing the growth of pathogens in food preservation, which may be relevant to their capability to secrete excessive organic acids into the environment to facilitate their dominance in the microbial community. Acetate metabolism is closely related to the regulation of pathogenic genes,^58^ which is only present in water and human groups, indicating that they may have similar pathogenic gene regulation patterns. Phenol 2-monooxygenase is only absent in the soil group of the *Ralstonia* genus, indicating that the phenol 2-monooxygenase associate genes were lost during adapted to the soil. Phthalates are widely used as plasticizers, and their natural degradation is slow, *R. pickettii* in the water and plant group possesses the potential pathway for biodegradation at the genomic level,^59^ which are strategies for survival using degraded organic matter. Malate dehydrogenase is an enzyme distributed in the central oxidative pathway,^60^ which is naturally present in Gram-negative bacteria as a dimeric molecule and is found in both the water and plant groups. PEP is one of the products of the reaction and can mediate carbohydrate uptake and phosphorylation, and it is involved in signal transduction.^61,62^ In this study, PEP production was found exclusively in groups associated with water and plant hosts, which may be linked to the requirement for invasion and adhesion in these hosts to produce specific proteins. The pyrimidine degradation pathway enables assimilation of nitrogen and carbon for growth,^59^ and *R. pickettii* in the water group employs this strategy, which may relate to its ability to survive in relatively oligotrophic media.^63^

### 4.4 The relationship between R. pickettii and Dolichospermum sp

Algae bloom is an important manifestation of microecological disorder in the water environment, manifested by an abnormal increase in the abundance of algae, extremely serious in China.^18^ *Dolichospermum* as one of bloom-forming cyanobacterium can produce toxins, including microcystin, which inhibit protein phosphatases, enhance cell membrane permeability, and cause DNA damage.^64-67^ These effects heighten the selection pressure for ARGs and facilitate HGT, potentially leading to a greater diversity of ARGs in aquatic environments of *R. pickettii*.^72^ We isolate *R. pickettii* from cyanobacterium cultures, enabling to preliminarily explore its possible relationship with algae blooms. Raising the question of whether its behavior similar to plant infestations, which adhere to the plant and then releases toxins. However, our findings show the different patterns between terrestrial plant parasitism and hydrophyte parasitism. The connection appears to be initially related to EPS include proteins, glyoxylates and lipids, which is critical for microbial aggregation^68-70^ and the microbial community can produce VB_12_ to promote the growth of cyanobacteria.^71^ In this study, due to *R. pickettii* having flagella and motility, it could potentially evade capture during experiment. The FISH results (Fig.5) indicate that *R. pickettii* may free living in lakes and achieve coexistence with cyanobacteria in a free state in the culture, they are attached around the algal filaments when nutrients are needed. Our study contributes to the understanding a *R. pickettii* reduced pathogenicity of water and opens an avenue for revealing that weather a reciprocal relationship between *R. pickettii* and *Dolichospermum* sp.

## 5. Conclusion

In conclusion, *Ralstonia* species are classified into four distinct groups based on habitat and host associations: soil, water, plant, and human. Analysis of ARGs reveals *OXA* and *ceoB* are the predominant types, with *ceoB* predominantly found in the plant-host associate clade and co-existence of *ceoB* and *OXA* in other clades. ARGs are especially categorized into two groups in those associated with human, underscoring the importance of distinguishing bacterial infections in clinical treatment. The annotation results of CAZymes highlight differences between plant group, water and human groups, with GHs mainly present in the plant group and GTs more abundant in other groups, correlating with host-specific infection patterns. The host specificity of *Ralstonia* spp. is reflected in the presence of the T3SS in the plant group but its incompleteness in other groups, particularly the water group, suggesting a reduced invasive capacity. The unique pyrimidine degradation pathway in water degrades exogenous pyrimidines to provide nitrogen and other substrates for energy metabolism. The results of secondary metabolism show that plant group associate is separated from other groups, this may play an important role in enhancing the competitiveness of endosymbiotic microbial communities in plants. These findings underscore the adaptive evolution of *Ralstonia* species across different habitats and hosts, accompanied by changes in virulence and invasion potential. During the period of algae blooms, the abundant *R. pickettii* camp free living and can use the EPS of algae to provide energy, the underlying relationship still needs further exploration.

## Data availability statement

Four strains of *R. pickettii* genomes supporting this study are deposited in the NCBI repository, accession numbers PRJNA1111332 (BioProject) and SAMN41388510, SAMN41388511, SAMN41388512, SAMN41388513 (BioSample).

## Author contributions

Investigation, Gaopeng Liu, Qi Li and Da Huo; Data Curation, Gaopeng Liu and Da Huo; Writing-Original Draft, Gaopeng Liu, Chengzhi Mao and Da Huo; Writing-Review & Editing, Gaopeng Liu, Qi Li, Chengzhi Mao and Da Huo; Supervision, Da Huo, Tao Li and Qi Li; Project Administration, Da Huo, Qi Li and Tao Li; Funding Acquisition, Da Huo and Tao Li;

## Acknowledgments

This study was supported by the National Key Research and Development Program of China (No. 2020YFA0907402), China Postdoctoral Science Foundation (No. 2021M703430), and the National Natural Science Foundation of China (No. 92251304). We thank Guangxin Wang (Analysis and Testing Center, IHB, CAS) for assistance with fluorescence microscopy scanning and analysis. Additionally, we would like to express our appreciation to Shengjie Sun (Tianjin Agricultural University) for his valuable contributions to the discussions.

## Limitations of the study

Due to the fact that this study is based on genome prediction results, additional direct experimental evidence is needed to support this interaction. Establish an indoor simulation group to observe the growth of *Dolichostermum* sp. and *R. pickettii* in coexistence, as well as the response to adding *R. pickettii* at different EPS concentrations. However, due to the difficulty in purifying *Dolichostermum*, indoor simulation experiments continue to encounter significant challenges.

## Reference

1. Ryan, M.P., Adley, C.C.J.E.j.o.c.m., and diseases, i. (2014). Ralstonia spp.: emerging global opportunistic pathogens. 33, 291–304.

2. Yuan, C., An, T., Li, X., Zou, J., Lin, Z., Gu, J., Hu, R., and Fang, Z.J.F.i.M. (2024). Genomic analysis of Ralstonia pickettii reveals the genetic features for potential pathogenicity and adaptive evolution in drinking water. 14, 1272636.

3. Fluit, A.C., Bayjanov, J.R., Aguilar, M.D., Cantón, R., Tunney, M.M., Elborn, J.S., van Westreenen, M., and Ekkelenkamp, M.B.J.A.V.L. (2021). Characterization of clinical Ralstonia strains and their taxonomic position. 114, 1721–1733.

4. Ryan, M., Pembroke, J., and Adley, C.J.J.o.H.i. (2006). Ralstonia pickettii: a persistent gram-negative nosocomial infectious organism. 62, 278–284.

5. Siddiqui, T., Patel, S.S., Sinha, R., Ghoshal, U., and Sahu, C.J.A.M. (2022). Ralstonia mannitolilytica: an emerging multidrug-resistant opportunistic pathogen in a tertiary care hospital setting. 4, 000367.

6. Stelzmueller, I., Biebl, M., Wiesmayr, S., Eller, M., Hoeller, E., Fille, M., Weiss, G., Lass-Floerl, C., Bonatti, H.J.C.M., and Infection (2006). Ralstonia pickettii—innocent bystander or a potential threat? 12(2),99–101.

7. Demirdag, T.B., Ozkaya-Parlakay, A., Bayrakdar, F., Gulhan, B., Yuksek, S.K., Yildiz, S.S., Mumcuoglu, İ., Dinc, B., Yarali, N.J.J.o.M., Immunology, and Infection (2022). An outbreak of Ralstonia pickettii bloodstream infection among pediatric leukemia patients. 55, 80–85.

8. Hayward, A.J.A.r.o.p. (1991). Biology and epidemiology of bacterial wilt caused by Pseudomonas solanacearum. 29, 65–87.

9. Qi, P., Huang, M., Hu, X., Zhang, Y., Wang, Y., Li, P., Chen, S., Zhang, D., Cao, S., and Zhu, W.J.T.P.C. (2022). A Ralstonia solanacearum effector targets TGA transcription factors to subvert salicylic acid signaling. 34, 1666–1683.

10. Peeters, N., Guidot, A., Vailleau, F., and Valls, M.J.M.p.p. (2013). R alstonia solanacearum, a widespread bacterial plant pathogen in the post-genomic era. 14, 651–662.

11. Chen, M.-Y., Teng, W.-K., Zhao, L., Hu, C.-X., Zhou, Y.-K., Han, B.-P., Song, L.-R., and Shu, W.-S.J.T.I.J. (2021). Comparative genomics reveals insights into cyanobacterial evolution and habitat adaptation. 15, 211–227.

12. Kumar, R., Verma, H., Haider, S., Bajaj, A., Sood, U., Ponnusamy, K., Nagar, S., Shakarad, M.N., Negi, R.K., and Singh, Y.J.M. (2017). Comparative genomic analysis reveals habitat-specific genes and regulatory hubs within the genus Novosphingobium. 2, 10.1128/msystems.00020-00017.

13. Wicker, E., Lefeuvre, P., De Cambiaire, J.-C., Lemaire, C., Poussier, S., and Prior, P.J.T.I.j. (2012). Contrasting recombination patterns and demographic histories of the plant pathogen Ralstonia solanacearum inferred from MLSA. 6, 961–974.

14. Safni, I., Cleenwerck, I., De Vos, P., Fegan, M., Sly, L., Kappler, U.J.I.j.o.s., and microbiology, e. (2014). Polyphasic taxonomic revision of the Ralstonia solanacearum species complex: proposal to emend the descriptions of Ralstonia solanacearum and Ralstonia syzygii and reclassify current R. syzygii strains as Ralstonia syzygii subsp. syzygii subsp. nov., R. solanacearum phylotype IV strains as Ralstonia syzygii subsp. indonesiensis subsp. nov., banana blood disease bacterium strains as Ralstonia syzygii subsp. celebesensis subsp. nov. and R. solanacearum phylotype I and III strains as Ralstonia pseudosolanacearum sp. nov. 64, 3087–3103.

15. Prior, P., Ailloud, F., Dalsing, B.L., Remenant, B., Sanchez, B., and Allen, C.J.B.g. (2016). Genomic and proteomic evidence supporting the division of the plant pathogen Ralstonia solanacearum into three species. 17, 1–11.

16. Lu, C.-H., Zhang, Y.-Y., Zhang, L.-Q., Jin, Y., and Xia, Z.-Y.J.F.i.M. (2023). Ralstonia chuxiongensis sp. nov., Ralstonia mojiangensis sp. nov., and Ralstonia soli sp. nov., isolated from tobacco fields, are three novel species in the family Burkholderiaceae. 14, 1179087.

17. Liu, Y., Wu, D., Liu, Q., Zhang, S., Tang, Y., Jiang, G., Li, S., and Ding, W.J.E.J.o.P.P. (2017). The sequevar distribution of Ralstonia solanacearum in tobacco-growing zones of China is structured by elevation. 147, 541–551.

18. Huo, D., Gan, N., Geng, R., Cao, Q., Song, L., Yu, G., and Li, R.J.H.A. (2021). Cyanobacterial blooms in China: Diversity, distribution, and cyanotoxins. 109, 102106.

19. Pritchard, L., Glover, R.H., Humphris, S., Elphinstone, J.G., and Toth, I.K.J.A.m. (2016). Genomics and taxonomy in diagnostics for food security: soft-rotting enterobacterial plant pathogens. 8, 12–24.

20. Schmieder, R., and Edwards, R.J.B. (2011). Quality control and preprocessing of metagenomic datasets. 27, 863–864.

21. Vaser, R., Sović, I., Nagarajan, N., and Šikić, M.J.G.r. (2017). Fast and accurate de novo genome assembly from long uncorrected reads. 27, 737–746.

22. Kolmogorov, M., Yuan, J., Lin, Y., and Pevzner, P.A.J.N.b. (2019). Assembly of long, error-prone reads using repeat graphs. 37, 540–546.

23. Hu, J., Fan, J., Sun, Z., and Liu, S.J.B. (2020). NextPolish: a fast and efficient genome polishing tool for long-read assembly. 36, 2253–2255.

24. Seemann, T.J.B. (2014). Prokka: rapid prokaryotic genome annotation. 30, 2068–2069.

25. Parks, D.H., Imelfort, M., Skennerton, C.T., Hugenholtz, P., and Tyson, G.W.J.G.r. (2015). CheckM: assessing the quality of microbial genomes recovered from isolates, single cells, and metagenomes. 25, 1043–1055.

26. Emms, D.M., and Kelly, S.J.G.b. (2019). OrthoFinder: phylogenetic orthology inference for comparative genomics. 20, 1–14.

27. Katoh, K., Misawa, K., Kuma, K.i., and Miyata, T.J.N.a.r. (2002). MAFFT: a novel method for rapid multiple sequence alignment based on fast Fourier transform. 30, 3059–3066.

28. Capella-Gutiérrez, S., Silla-Martínez, J.M., and Gabaldón, T.J.B. (2009). trimAl: a tool for automated alignment trimming in large-scale phylogenetic analyses. 25, 1972–1973.

29. Shen, W., Le, S., Li, Y., and Hu, F.J.P.o. (2016). SeqKit: a cross-platform and ultrafast toolkit for FASTA/Q file manipulation. 11, e0163962.

30. Nguyen, L.-T., Schmidt, H.A., Von Haeseler, A., Minh, B.Q.J.M.b., and evolution (2015). IQ-TREE: a fast and effective stochastic algorithm for estimating maximum-likelihood phylogenies. 32, 268–274.

31. Letunic, I., and Bork, P.J.N.a.r. (2021). Interactive Tree Of Life (iTOL) v5: an online tool for phylogenetic tree display and annotation. 49, W293-W296.

32. Aramaki, T., Blanc-Mathieu, R., Endo, H., Ohkubo, K., Kanehisa, M., Goto, S., and Ogata, H.J.B. (2020). KofamKOALA: KEGG Ortholog assignment based on profile HMM and adaptive score threshold. 36, 2251–2252.

33. Zhang, H., Yohe, T., Huang, L., Entwistle, S., Wu, P., Yang, Z., Busk, P.K., Xu, Y., and Yin, Y.J.N.a.r. (2018). dbCAN2: a meta server for automated carbohydrate-active enzyme annotation. 46, W95-W101.

34. Zankari, E., Hasman, H., Cosentino, S., Vestergaard, M., Rasmussen, S., Lund, O., Aarestrup, F.M., and Larsen, M.V.J.J.o.a.c. (2012). Identification of acquired antimicrobial resistance genes. 67, 2640–2644.

35. Chen, L., Yang, J., Yu, J., Yao, Z., Sun, L., Shen, Y., and Jin, Q.J.N.a.r. (2005). VFDB: a reference database for bacterial virulence factors. 33, D325–D328.

36. Blin, K., Shaw, S., Augustijn, H.E., Reitz, Z.L., Biermann, F., Alanjary, M., Fetter, A., Terlouw, B.R., Metcalf, W.W., and Helfrich, E.J.J.N.a.r. (2023). antiSMASH 7.0: new and improved predictions for detection, regulation, chemical structures and visualisation. 51, W46-W50.

37. Librado, P., Vieira, F.G., and Rozas, J.J.B. (2012). BadiRate: estimating family turnover rates by likelihood-based methods. 28, 279–281.

38. Delsuc, F., Brinkmann, H., and Philippe, H.J.N.R.G. (2005). Phylogenomics and the reconstruction of the tree of life. 6, 361–375.

39. Newman, D.J., Cragg, G.M., and Snader, K.M.J.J.o.n.p. (2003). Natural products as sources of new drugs over the period 1981− 2002. 66, 1022-1037.

40. Nordmann, P., Poirel, L., Kubina, M., Casetta, A., Naas, T.J.A.a., and chemotherapy (2000). Biochemical-genetic characterization and distribution of OXA-22, a chromosomal and inducible class D β-lactamase from Ralstonia (Pseudomonas) pickettii. 44, 2201-2204.

41. Guglierame, P., Pasca, M.R., De Rossi, E., Buroni, S., Arrigo, P., Manina, G., and Riccardi, G.J.B.m. (2006). Efflux pump genes of the resistance-nodulation-division family in Burkholderia cenocepacia genome. 6, 1–14.

42. Bellanger, X., Guilloteau, H., Bonot, S., and Merlin, C.J.S.o.t.T.E. (2014). Demonstrating plasmid-based horizontal gene transfer in complex environmental matrices: a practical approach for a critical review. 493, 872–882.

43. Gootz, T.D.J.C.R.i.I. (2010). The global problem of antibiotic resistance. 30(1).

44. Costa, T.R., Felisberto-Rodrigues, C., Meir, A., Prevost, M.S., Redzej, A., Trokter, M., and Waksman, G.J.N.R.M. (2015). Secretion systems in Gram-negative bacteria: structural and mechanistic insights. 13, 343–359.

45. Poueymiro, M., and Genin, S.J.C.o.i.m. (2009). Secreted proteins from Ralstonia solanacearum: a hundred tricks to kill a plant. 12, 44–52.

46. Sandkvist, M., Michel, L.O., Hough, L.P., Morales, V.M., Bagdasarian, M., Koomey, M., DiRita, V.J., and Bagdasarian, M.J.J.o.b. (1997). General secretion pathway (eps) genes required for toxin secretion and outer membrane biogenesis in Vibrio cholerae. 179, 6994–7003.

47. Xia, X.-h., Liu, G.-p., Wu, X.-l., Cui, S.-s., Yang, C.-H., Du, Q.-y., and Zhang, X.-w.J.A. (2021). Effects of Macleaya cordata extract on TLR20 and the proinflammatory cytokines in acute spleen injury of loach (Misgurnus anguillicaudatus) against Aeromonas hydrophila infection. 544, 737105.

48. Asolkar, T., and Ramesh, R.J.J.o.g. (2018). Identification of virulence factors and type III effectors of phylotype I, Indian Ralstonia solanacearum strains Rs-09-161 and Rs-10-244. 97, 55-66.

49. Toth, I.K., and Birch, P.R.J.C.o.i.p.b. (2005). Rotting softly and stealthily. 8, 424–429.

50. Huang, L., Zhang, H., Wu, P., Entwistle, S., Li, X., Yohe, T., Yi, H., Yang, Z., and Yin, Y.J.N.A.R. (2018). dbCAN-seq: a database of carbohydrate-active enzyme (CAZyme) sequence and annotation. 46, D516-D521.

51. Poole, J., Day, C.J., von Itzstein, M., Paton, J.C., and Jennings, M.P.J.N.R.M. (2018). Glycointeractions in bacterial pathogenesis. 16, 440–452.

52. Szymanski, C.M., and Wren, B.W.J.N.R.M. (2005). Protein glycosylation in bacterial mucosal pathogens. 3, 225–237.

53. Lu, Q., Li, S., and Shao, F.J.T.i.m. (2015). Sweet talk: protein glycosylation in bacterial interaction with the host. 23, 630–641.

54. Dang, T., and Su ssmuth, R.D.J.A.o.c.r. (2017). Bioactive peptide natural products as lead structures for medicinal use. 50, 1566–1576.

55. Somayaji, A., Dhanjal, C.R., Lingamsetty, R., Vinayagam, R., Selvaraj, R., Varadavenkatesan, T., and Govarthanan, M.J.M.R. (2022). An insight into the mechanisms of homeostasis in extremophiles. 263, 127115.

56. Tymiak, A.A., Culver, C.A., Malley, M.F., and Gougoutas, J.Z.J.T.J.o.O.C. (1985). Structure of obafluorin: an antibacterial. beta.-lactone from Pseudomonas fluorescens. 50, 5491–5495.

57. Pinhal, S., Ropers, D., Geiselmann, J., and De Jong, H.J.J.o.b. (2019). Acetate metabolism and the inhibition of bacterial growth by acetate. 201, 10.1128/jb.00147-00119.

58. Bernal, V., Castaño-Cerezo, S., Cánovas, M.J.A.m., and biotechnology (2016). Acetate metabolism regulation in Escherichia coli: carbon overflow, pathogenicity, and beyond. 100, 8985–9001.

59. Yin, J., Wei, Y., Liu, D., Hu, Y., Lu, Q., Ang, E.L., Zhao, H., and Zhang, Y.J.J.o.B.C. (2019). An extended bacterial reductive pyrimidine degradation pathway that enables nitrogen release from β-alanine. 294, 15662–15671.

60. Takahashi-Íñiguez, T., Aburto-Rodríguez, N., Vilchis-González, A.L., and Flores, M.E.J.J.o.Z.U.-S.B. (2016). Function, kinetic properties, crystallization, and regulation of microbial malate dehydrogenase. 17, 247–261.

61. Postma, P.W., Lengeler, J.W., and Jacobson, G.J.M.r. (1993). Phosphoenolpyruvate: carbohydrate phosphotransferase systems of bacteria. 57, 543–594.

62. Sun, S., Qiao, Z., Sun, K., and Huo, D.J.M.S. (2024). Assembly process and co-occurrence network of microbial community in response to free ammonia gradient distribution. e01051–01024.

63. Matache, R., Deak, G., Jawdhari, A., Sadica, I., Pop, C.-E., Fendrihan, S., and Craciun, N.J.b. (2024). First insights of the Danube sturgeon (Acipenser gueldenstaedtii) skin adherent microbiota. 2024.2003. 2013.584882.

64. Ji, W., Ma, J., Zheng, Z., Al-Herrawy, A.Z., Xie, B., and Wu, D.J.S.o.T.T.E. (2024). Algae blooms with resistance in fresh water: Potential interplay between Microcystis and antibiotic resistance genes. 173528.

65. Hitzfeld, B.C., Höger, S.J., and Dietrich, D.R.J.E.h.p. (2000). Cyanobacterial toxins: removal during drinking water treatment, and human risk assessment. 108, 113–122.

66. Jong, M.-C., Harwood, C.R., Blackburn, A., Snape, J.R., Graham, D.W.J.E.S., and Technology (2020). Impact of redox conditions on antibiotic resistance conjugative gene transfer frequency and plasmid fate in wastewater ecosystems. 54, 14984–14993.

67. Winter, M., Buckling, A., Harms, K., Johnsen, P.J., and Vos, M.J.C.O.i.M. (2021). Antimicrobial resistance acquisition via natural transformation: context is everything. 64, 133–138.

68. Liu, Y., Yang, F., Liu, S., Zhang, X., and Li, M.J.S.o.T.T.E. (2023). Molecular characteristics of microalgal extracellular polymeric substances were different among phyla and correlated with the extracellular persistent free radicals. 857, 159704.

69. Naveed, S., Li, C., Lu, X., Chen, S., Yin, B., Zhang, C., Ge, Y.J.C.R.i.E.S., and Technology (2019). Microalgal extracellular polymeric substances and their interactions with metal (loid) s: A review. 49, 1769–1802.

70. Guan, Y., Yu, G., Jia, N., Han, R., and Huo, D.J.E.I. (2024). Spectral characteristics of dissolved organic matter in Plateau Lakes: Identifying eutrophication indicators in Southwest China. 82, 102703.

71. Xie, M., Ren, M., Yang, C., Yi, H., Li, Z., Li, T., and Zhao, J.J.F.i.M. (2016). Metagenomic analysis reveals symbiotic relationship among bacteria in Microcystis-dominated community. 7, 173284.

