## Supplementary figures and images for "Comparative genomic analysis reveals reduced pathogenicity of *Ralstonia* spp. in water"

### Supplemental Figure 1

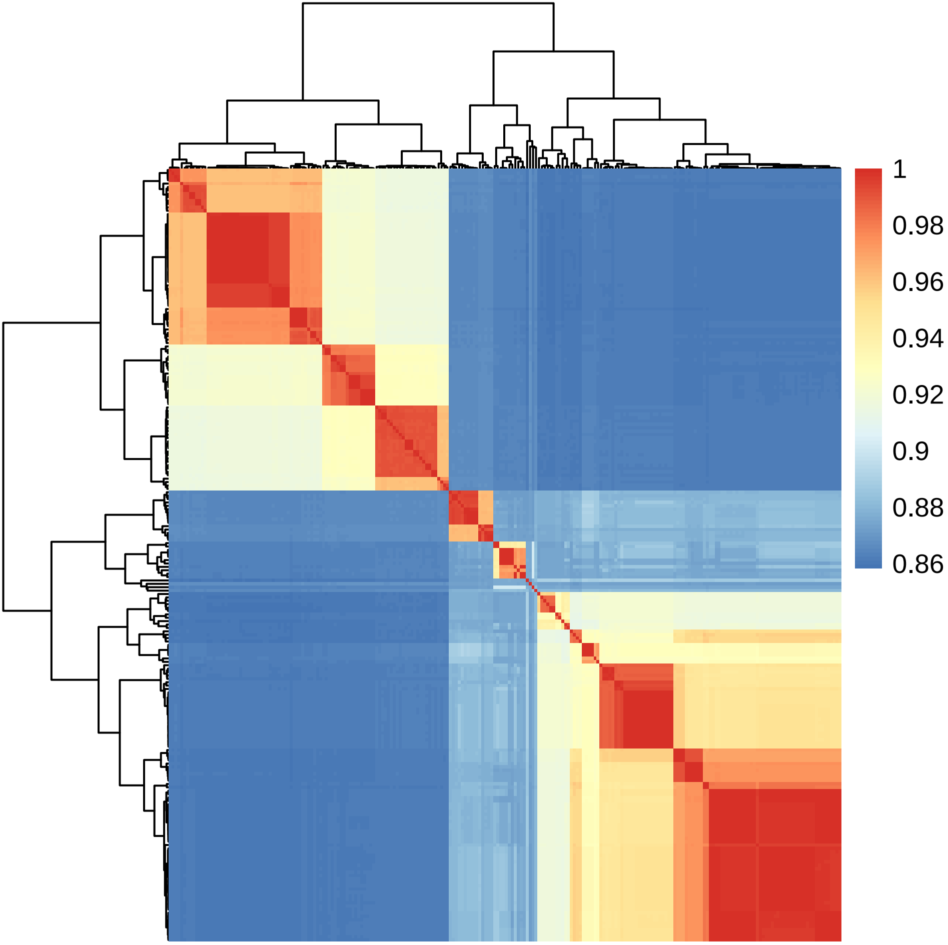

### Supplemental Figure 2

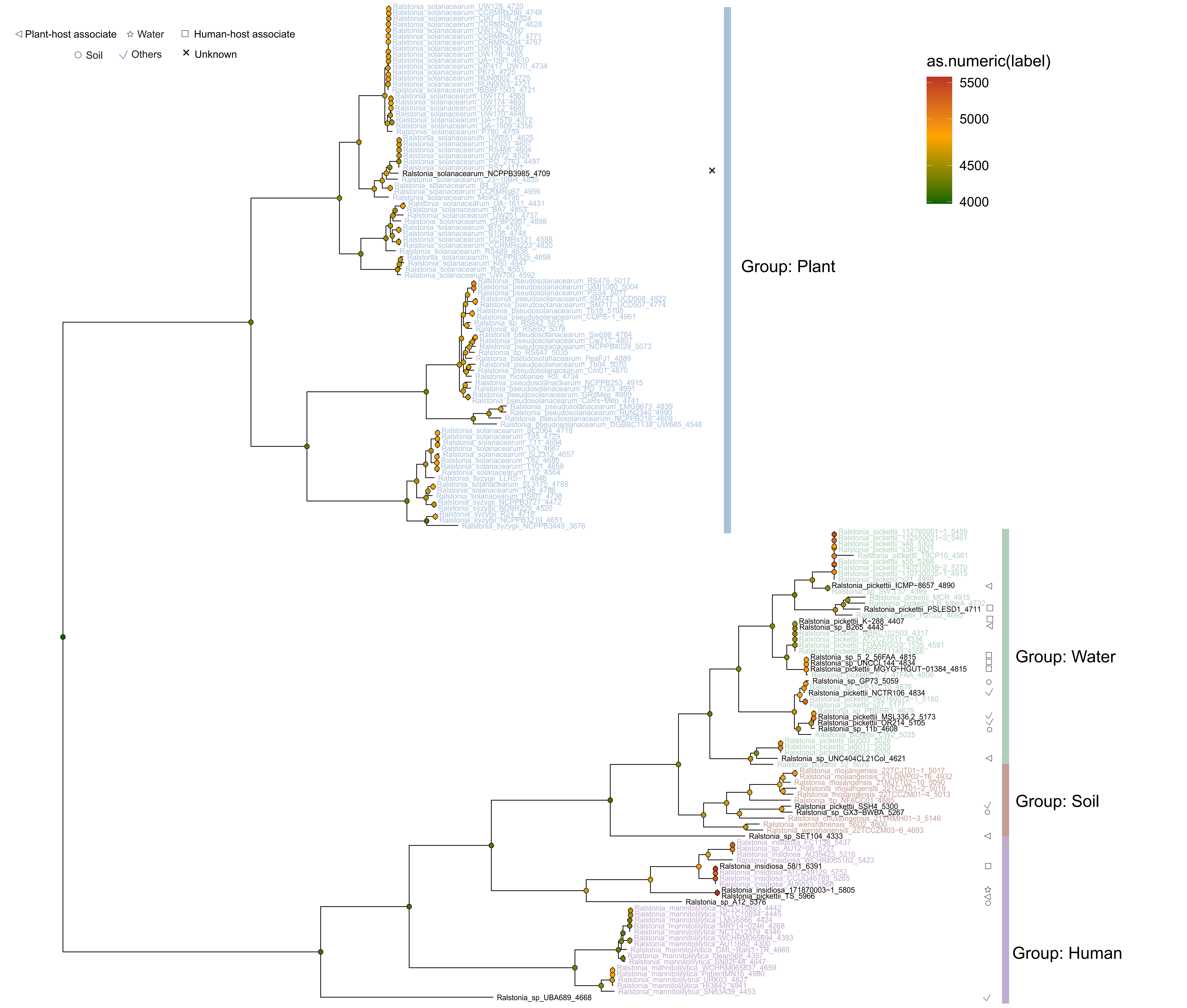
